# Transcriptome and miRNAome reveal components regulating primary thickening of bamboo shoots

**DOI:** 10.1101/2021.09.23.461506

**Authors:** Ying Li, Deqiang Zhang, Yongfeng Lou, An Xinmin, Zhimin Gao

**Affiliations:** National State Forestry and Grassland Administration Key Open Laboratory on the Science and Technology of Bamboo and Rattan, Institute of Gene Science for Bamboo and Rattan Resources, International Centre for Bamboo and Rattan, Beijing 100102, China; National Engineering Laboratory for Tree Breeding, College of Biological Sciences and Technology, Beijing 100083, China; Jiangxi Academy of Forestry, Jiangxi 330032, China

**Author notes:** Y. L. conceived the original screening and research plans; designed, performed the experiments and analyzed the data; wrote the article with contributions of all the authors; DQ. Z. designed the experiments and supervised the writing; YF. L. and XM. A. provided technical assistance to Y. L.; ZM. G. supervised the experiments and the writing, and agrees to serve as the author responsible for contact and ensures communication. (Y. L.); (DQ. Z.); (YF. L.); (XM. A.); (ZM. G.).

## Abstract

Primary thickening determines bamboo yield and wood property. However, little is known about the regulatory networks involved in this process. The present study identified a total of 58,652 genes and 521 miRNAs via transcriptome and small RNA sequencing using the underground thickening shoot samples of wild type (WT) Moso bamboo (*Phyllostachys edulis*) and a thick wall (TW) variant (*P. edulis* cv. *Pachyloen*) at five developmental stages (WTS1/TWS1-WTS5/TWS5). A total of 11,636 (54.05%) differentially expressed genes (DEGs) and 515 (98.85%) differentially expressed miRNAs (DEmiRs) were identified from the WT, TW, and WTTW groups. The first two groups were composed of four pairwise combinations each between two successive stages (WTS2/TWS2_vs_WTS1/TWS1, WTS3/TWS3_vs_WTS2/TWS2, WTS4/TWS4_vs_WTS3/TWS3 and WTS5/TWS5_vs_WTS4/TWS4), and the WTTW group was composed of five between two relative stages (TWS1–5_vs_WTS1–5). Additionally, among the phytohormones, zeatin (ZT) showed more remarkable changes in concentrations than indole-3-acetic acid (IAA), gibberellic acid (GA_3_), and abscisic acid (ABA) throughout the five stages in the WT and the TW groups. Moreover, 118 sites were identified for 590 miRNA-mRNA pairs via degradome sequencing. The dual-luciferase reporter assay confirmed that 14 miRNAs bound to 12 targets. Fluorescence in situ hybridization (FISH) localized miR166 and miR160 in the shoot apical meristem (SAM) and the procambium of Moso bamboo shoots at the S1 stage. Thus, primary thickening is a complex process regulated by miRNA-gene-phytohormone networks, and the miRNAome and transcriptome dynamics regulate phenotypic plasticity. These findings provide insights into the molecular mechanisms underlying wood formation and properties and propose targets for bamboo breeding.

## Introduction

Bamboo plants are major wood resources, which belong to the Bambusoideae subfamily of the Poaceae grass family. They are characterized by rapid growth and excellent wood quality like perennial woody dicots and have well-developed rhizomes, with buds at the nodes that develop into aboveground hollow culms. However, the culms do not show secondary growth due to the lack of interfascicular cambium. Interestingly, the shoots of Moso bamboo (*Phyllostachys edulis* (Carr.) H. de Lehaie), an economically important species known for its taste and nutritional value, exhibit underground primary thickening during which the bud gradually develops into a mature shoot. Subsequently, the mature shoot grows fast, with the elongation of aboveground internodes maintaining a constant node number and coarseness. Primary thickening largely determines the morphological features and the anatomical characteristics of bamboo mature culms. Therefore, a better understanding of the molecular regulation of the primary growth is essential for improving bamboo biomass and wood property and the shoots edible value.

Shoot apical meristem (SAM) maintenance and organogenesis are responsible for primary thickening, accompanied by various physiological and biochemical changes, including carbohydrate metabolism and energy transformation. Many genes have been identified involved in plant hormone signal transduction and cell wall development (Wei et al., 2017) identified in bamboo shoots. Genes related to programmed cell death, ethylene and calcium signaling pathways, protein hydrolysis, and nutrient cycle have also been identified in *Pseudosasa japonica* (Guo et al., 2018). Meanwhile, Li et al. (2018) detected changes in hormone pathways in the SAM during rapid aboveground internode elongation in Moso bamboo. Recently, Wang et al. (2021) identified miRNAs in the aboveground internodes of Moso bamboo’s mature culms, suggesting their role in rapid internode elongation. Although bamboo research has gradually progressed, studies are limited to functional genes identification.

In plants, miRNAs guide the Argonaute (AGO)-centered RNA-induced silencing complexes (RISCs) to target genes (Rodriguez et al., 2016). For example, miR165 and miR166 target the homeodomain-leucine zipper (HD-ZIP) transcription factor (TF) family to regulate meristem development and organ polarity (Byrne, 2006; Zhou et al., 2007). Further studies established that AGO10 inhibits miR165/166 activity, leading to a localized enrichment of HD-ZIP transcripts for programming SAM from the developing embryos to the entire adaxial domain and vasculature of cotyledons and new leaf primordia (Zhang and Zhang, 2012; Zhou et al., 2015). Meanwhile, expression profile analysis in soybean identified the putative role of miRNA166 and miRNA390 in SAM and bud development (Wong et al., 2011). Researchers have also identified miRNAs in bamboo, especially in aboveground parts, such as flowers (Ge et al., 2017). However, little is known on the role of miRNAs and the regulatory network in the primary thickening bamboo shoots.

The miRNAs are relatively conservative, while a few demonstrate species-specific regulatory effects and mechanisms. For example, *Arabidopsis* miR156 targets specific members of the SPL (SQUAMOSA promoter binding protein-like 14) TF family, promoting the transition from vegetative to reproductive phase (Wu and Poethig, 2006). In rice, miR156 affected tillering by regulating *OsSPL14* expression (Luo et al., 2012). Furthermore, changes in the expression levels of miRNAs of the same family have been observed in plants of the same family and genus, indicating differences in regulatory effects. The response patterns of miRNAs in Euphorbiaceae plants under abiotic stress showed their species- and stress-specific functions (Zeng et al., 2010). D’Ario (2017) suggested the role of miRNAs in establishing plant cell identity and creating diverse plant organ forms and shapes during plant evolution. Therefore, identifying the conserved and novel miRNAs will provide new insights into the molecular mechanism underlying primary thickening and morphological diversity in bamboo.

The thick wall (TW) stable moso bamboo variant (*P. edulis* cv. Pachyloen) has a narrow pith cavity, increased vascular bundles, and a fasciated culm (Guo, 2003; Guo et al., 2005). However, the molecular mechanism underlying the peculiar phenotype remains unclear. The present study aimed to (1) detect the key miRNAs and/or regulatory genes and delineate the interplay, and (2) discover differences in transcriptome and miRNAome between the wild-type (WT) and the TW variant and assess their roles in promoting morphological novelty. A comprehensive and integrated analysis was performed using thickening bamboo shoots at five different stages collected from the WT Moso bamboo and the TW variant (Figure 1). The study’s findings will provide new insights into the regulatory components and transcriptome dynamics during primary thickening and propose candidates for biomass and wood property improvement in bamboo.

**Figure 1.**
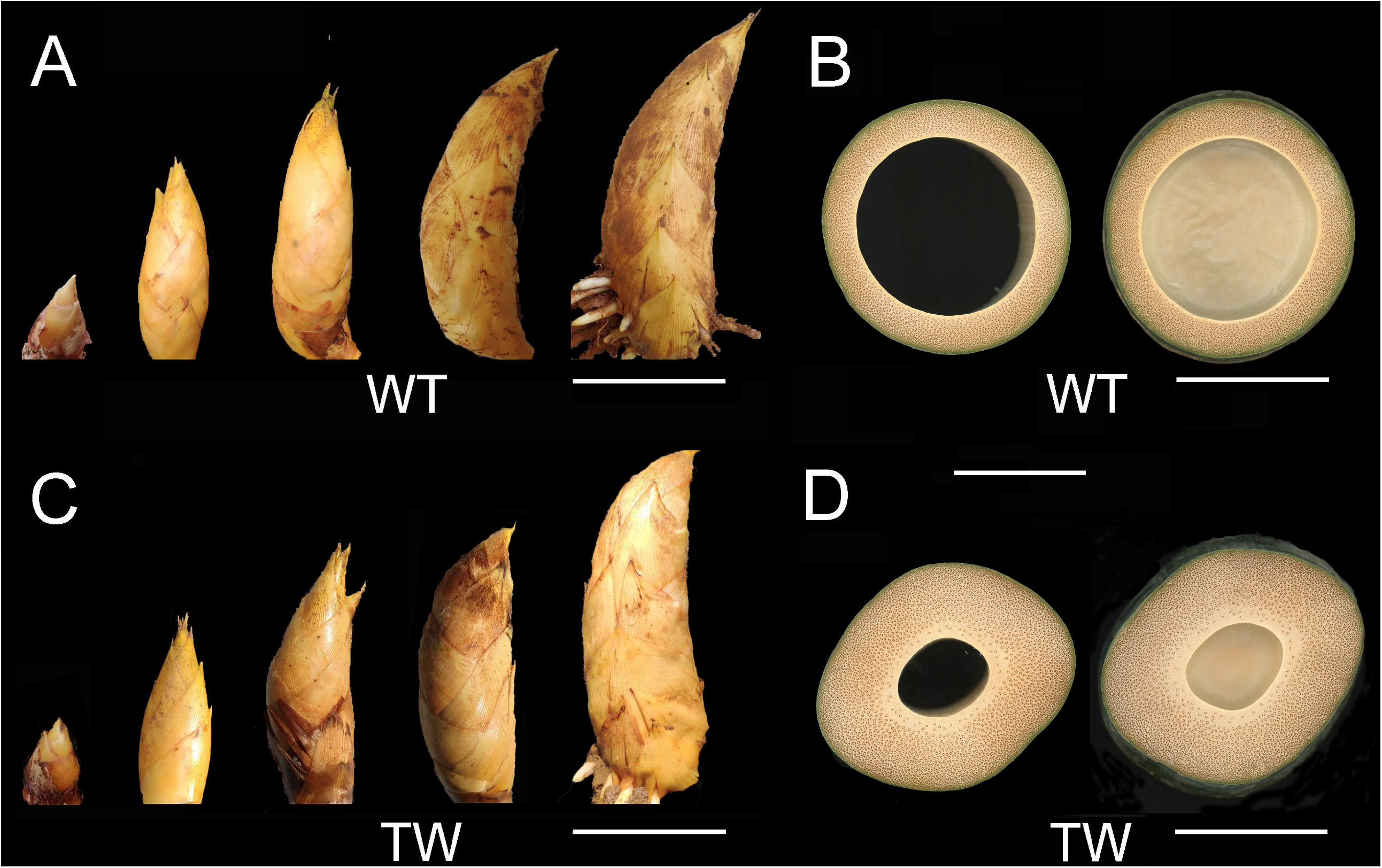
Aboveground mature culms and underground shoots of the wild-type (WT) moso bamboo and the thick wall (TW) variant. Underground shoots collected at five different stages (S1-S5), and cross-sections of the third intranode and internode of the equal-diameter aboveground mature culms of the WT Moso bamboo (A and B) and the TW variant (C and D). Bar = 10 cm.

## Results

### Transcriptome sequencing identified 7716 novel genes

Transcriptome sequencing of ten samples, including the WT and the TW variant at five developmental stages (WT1–WT5 and TW1–TW5), generated a total of 221.76 Gb high-quality reads (Q30 > 92.98%), with at least 20.08 Gb from each sample. The reads were uniquely aligned with the Moso bamboo reference genome v2.0 (Zhao et al., 2018) at an efficiency ranging from 91.20%–92.10% (Table 1). The reference-guided assembly of the mapped reads using the Cufflinks/Cuffmerge pipeline identified a total of 58,652 genes, of which 7,716 (13.16%) were novel ones. A total of 21,527 genes were expressed (FPKM ≥ 1) in at least one of the samples.

**Table 1.** Summary of transcriptome and small RNA sequencing data generated for 10 samples.

| Sample | Transcriptome sequencing |  |  | Small RNA sequencing |  |  |  |  |
| --- | --- | --- | --- | --- | --- | --- | --- | --- |
| | No. of clean reads<br>(single-ended)<sup>s</sup> | %≥Q<br>30 (%)<sup>a</sup> | No. of uniquely mapped<br>reads (%)<sup>s</sup> | No. of clean reads<br>(%)<sup>s</sup> | Q30 (%)<sup>a</sup> | GC<br>content<sup>a</sup> | No. of unannotated<br>reads<sup>s</sup> | No. of mapped<br>reads (%)<sup>s</sup> |
| TW1 | 141,213,840 | 93.10 | 129,479,444 (91.69) | 38390267 (89.00) | 98.70 | 53.67 | 6827998 | 4239380 (62.10) |
| TW2 | 140,389,470 | 93.86 | 128,029,273 (91.20) | 49565551 (87.83) | 98.77 | 51.97 | 12852242 | 8652591 (67.71) |
| TW3 | 140,521,926 | 93.21 | 128,163,980 (91.21) | 50650073 (93.49) | 98.82 | 50.78 | 13474182 | 9658538 (71.55) |
| TW4 | 137,370,524 | 93.48 | 125,872,244 (91.63) | 54045389 (92.60) | 98.49 | 51.58 | 13715823 | 9231245 (66.93) |
| TW5 | 166,876,286 | 93.45 | 152,712,950 (91.51) | 42701803 (76.62) | 98.68 | 51.06 | 6413479 | 4333974 (63.85) |
| WT1 | 141,213,840 | 92.98 | 129,479,444 (91.69) | 36908724 (82.07) | 99.03 | 52.88 | 6745208 | 3941662 (58.20) |
| WT2 | 156,284,680 | 93.22 | 142,866,399 (91.41) | 39072098 (89.00) | 99.09 | 52.09 | 9735327 | 6460195 (66.27) |
| WT3 | 149,074,996 | 93.36 | 136,621,922 (91.64) | 38731366 (90.69) | 99.06 | 51.94 | 10150773 | 7024012 (69.30) |
| WT4 | 134,363,964 | 93.65 | 123,399,863 (91.84) | 54571591 (90.37) | 98.62 | 51.80 | 13442651 | 9394293 (70.03) |
| WT5 | 167,376,092 | 93.63 | 154,158,971 (92.10) | 47133901 (91.19) | 98.90 | 52.34 | 11169855 | 7819337 (69.97) |

s: sum of data collected from three biological replications;
a: average of data;
Uniquely mapped reads: Reads that were uniquely mapped onto the reference genome;
Unannotated reads: reads used for alignment onto the reference genomes.

### Genotype- and/or stage-specific expression of genes

Furthermore, the differentially expressed genes (DEGs) were identified from 13 pairwise combinations of the WT, TW, and WTTW groups. The first two groups were composed of four pairwise combinations each between two successive stages (WTS2/TWS2_vs_WTS1/TWS1, WTS3/TWS3_vs_WTS2/TWS2, WTS4/TWS4_vs_WTS3/TWS3 and WTS5/TWS5_vs_WTS4/TWS4), and the WTTW group was composed of five between two relative stages (TWS1–5_vs_WTS1–5). A total of 11,636 (54.05%) DEGs were identified from the 13 pairwise combinations, with the number ranging from 131 in S2 of the WTTW group to 5,351 in S2_vs_S1 of the WTTW group (Figure 2A). These observations indicate the genotype- and/or stage-specific differential expression of genes. The TW group had the maximum DEGs (9,026), followed by the WTTW group (5,565), while the WT group had the minimum (3,177). Interestingly, S2_vs_S1 had the maximum DEGs for both the WT (1,815) and the TW (5,351) groups. Many DEGs were also identified in WTS4_vs_WTS3 (1,235) and TWS3_vs_TWS2 (4,942). In the WTTW group, the maximum DEGs were identified at S3 (4,721).

**Figure 2.**
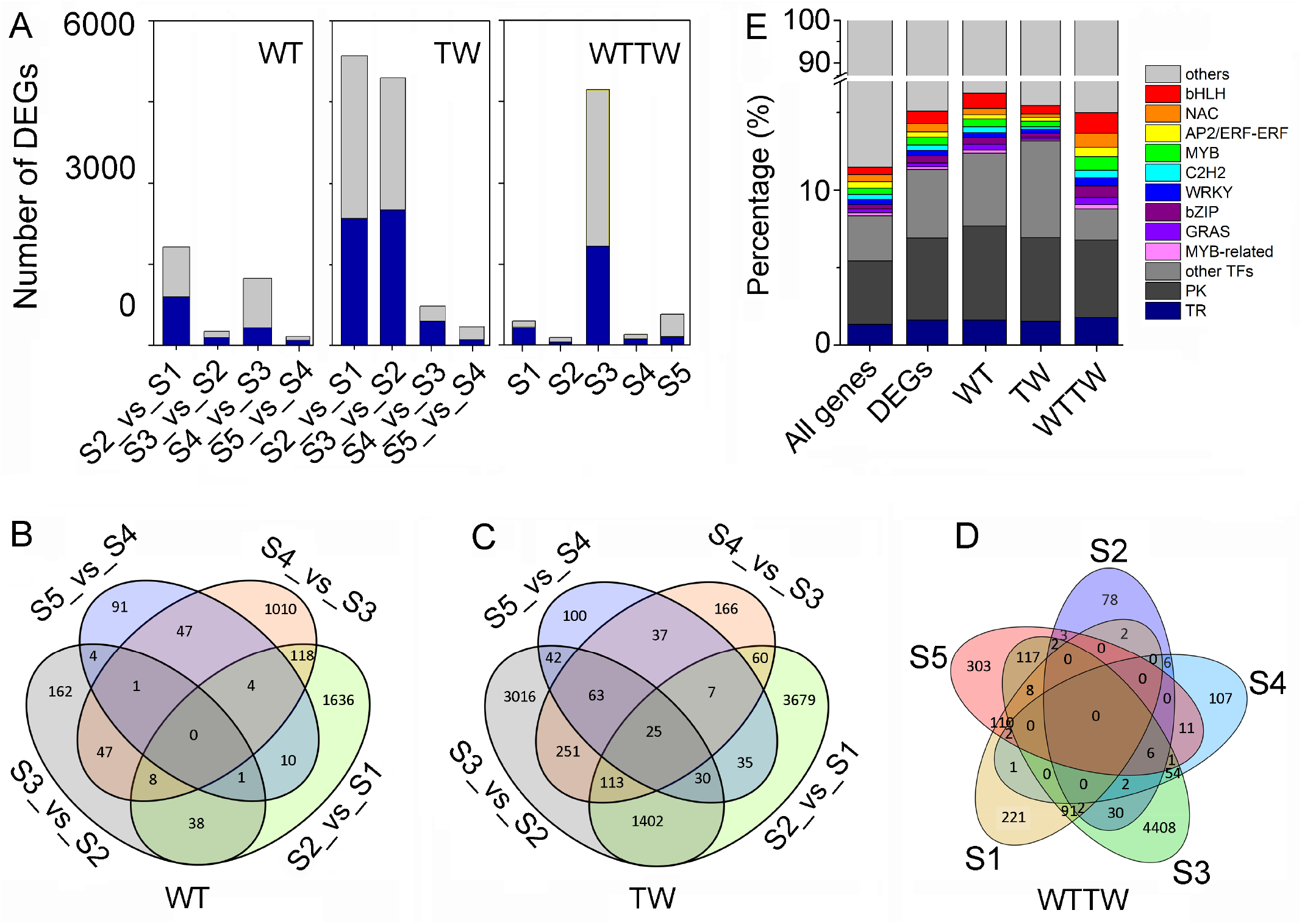
Differentially expressed genes (DEGs) identified from 13 pairwise combinations of the WT, TW, and WTTW groups. Both the wild-type (WT) and the thick wall (TW) groups were composed of four pairwise combinations between two successive stages (WTS2/TWS2_vs_WTS1/TWS1, WTS3/TWS3_vs_WTS2/TWS2, WTS4/TWS4_vs_WTS3/TWS3, and WTS5/TWS5_vs_WTS4/TWS4), and the WTTW group was composed of five pairwise combinations between two relative stages (TWS1– 5_vs_WTS1–5). A, DEGs ranged from 131 in S2 of the WTTW group to 5,351 in S2_vs_S1 of the WTTW group, with the upregulated (light grey) and downregulated (dark blue) genes almost equally representing few pairwise combinations. B–D, A total of 2899, 6961, and 5117 DEGs showed stage- and/or genotype-specific differential expression in the WT (B), TW (C), and WTTW (D) groups. E, The percentage of differentially expressed transcription factor (TF), phosphokinase (PK), and transcription regulator (TR) genes among the total genes identified, total DEGs, and DEGs in the WT, TW, and WTTW groups.

Furthermore, 2,899 (91.25%) and 6,961 (77.12%) DEGs showed stage-specific differential expression in the WT and TW groups, respectively, while 5,117 (91.95%) exhibited genotype-specific differential expression in the WTTW group (Figure 2B–D). Meanwhile, 25 DEGs were found in all four pairwise combinations of the TW group and none in either WT or WTTW group.

### DEGs of different pairwise combinations shared the GO terms

A total of 2,265 (71.29%), 6,414 (71.06%), and 3,930 (70.62%) DEGs were assigned GO terms in WT, TW, and WTTW groups, respectively. As shown in Figure S1 and Table S1, DEGs identified from various different pairwaise combinations shared several GO terms under the biological process category, including “DNA replication initiation (GO:0006270)”, “regulation of transcription, DNA-templated (GO:0006355)”, “auxin efflux (GO:0010315)”, and “microtubule-based movement (GO:0007018)”. The KEGG analysis indicated that DEGs in the WT, TW, and WTTW groups were enriched in 116, 125, and 123 pathways, respectively; DNA replication (ko03030) was the shared pathway among these three groups (Figure S2).

### Several genes encoding transcription factors (TFs) showed differential expression

In this study, a total of 3,543 genes, including 3,496 known and 47 novel genes encoding TFs belonging to 66 families, were identified. The maximum TF families included bHLH (basic helix-loop-helix; 270, 7.62%), NAC (NAM, ATAF1/2, and CUC2; 267, 7.54%), AP2/ERF (APETALA2/Ethylene responsive factor; 246, 6.94%) and MYB (v-myb avian myeloblastosis viral oncogene homolog, 242, 6.83%) (Figure 2E). In addition, 2,397 and 780 genes encoding protein kinases (PKs) and transcription regulators (TRs), respectively, were identified, of which 617 PKs and 188 TRs showed differential expression between the successive and relative stages of the WT, the TW and the WTTW groups (Table S2).

The differentially expressed TF genes accounted for a more significant percentage (8.17%) of total DEGs than the PK (5.30%) and TR (1.62%) genes, while their proportions were similar in the WT (8.37%, 6.07%, and 1.61%), TW (8.40%, 5.38%, and 1.54%), and WTTW (7.89%, 5.00%, and 1.80%) groups. Besides, the percentage of the nine most abundant TF families was roughly the same in the three groups.

### Profiles of DEGs were similar between WT and TW across five stages, except for S3

Clustering analysis showed that 11,608 (99.76%) DEGs out of 11,636 formed 11 clusters (clusters I–XI), divided into four groups based on the expression pattern under the five stages (Figure 3A). Interestingly, DEGs in most groups bend in the opposite direction at S3 in both WT and TW, although their trends were nearly the same; a few GO terms, including “regulation of transcription, DNA-templated” and “proteasome-mediated ubiquitin-dependent protein catabolic process (GO:0043161)” were shared among the clusters.

**Figure 3.**
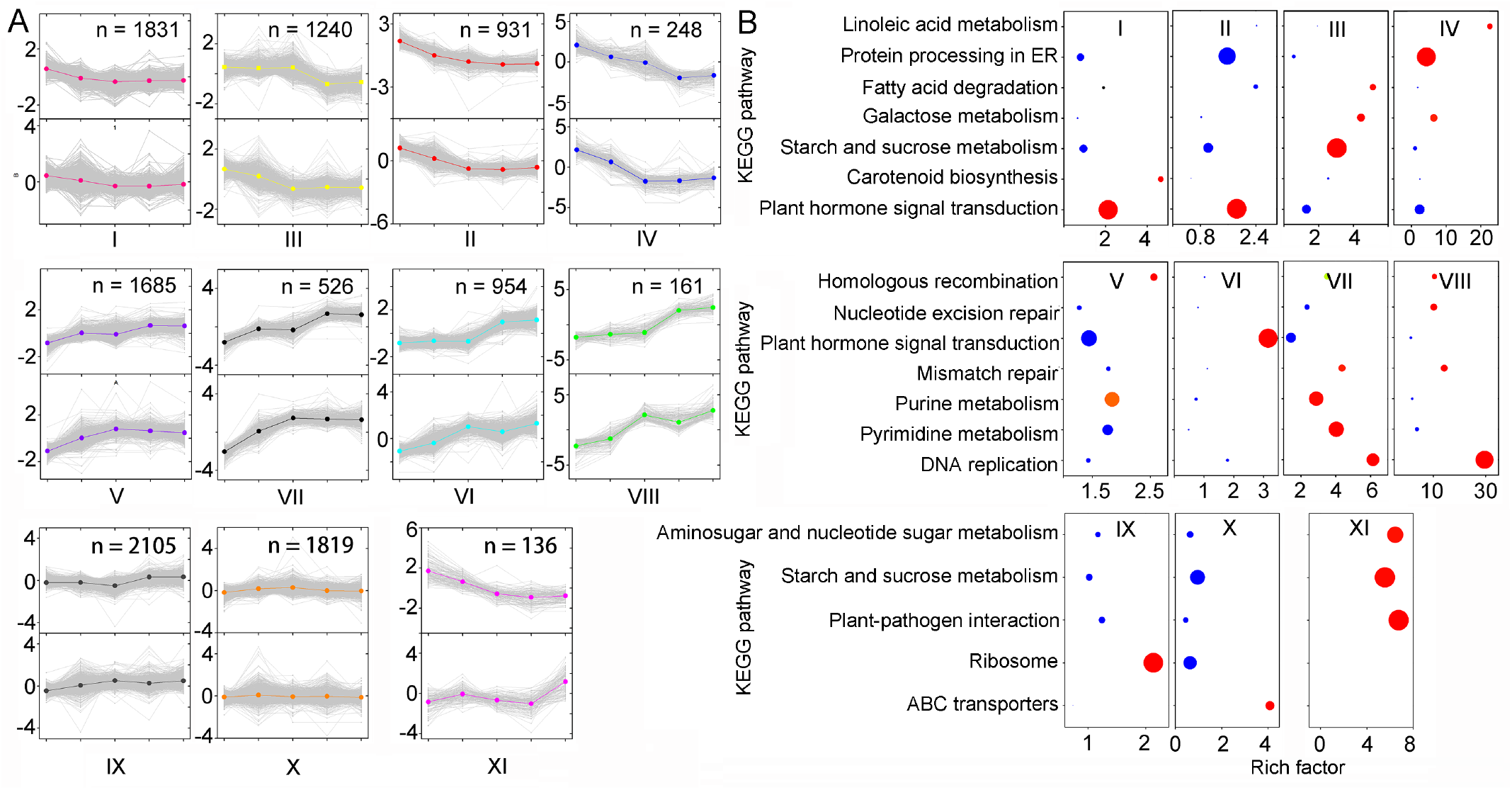
Clustering and KEGG pathway enrichment analysis of differentially expressed genes (DEGs) in the wild-type (WT) and the thick wall (TW) groups. A-B, Clustering and KEGG pathway enrichment analysis of DEGs across five unique stages (S1–S5) in the WT (the upper half) and the TW (the second half) groups. ER = endoplasmic reticulum. The size of dots in Figure 3B is proportional to the number of genes.

Several GO terms and KEGG pathways of the DEGs were common among the different clusters (KS < 0.005, FDR < 0.05; Figure 3B and Table S3, S4). Clusters I, II, and VI shared the significantly enriched GO term “intracellular signal transduction (GO:0035556)” and the KEGG pathway plant hormone signal transduction (ko04075); clusters III and IV shared “glutamine metabolic process (GO:0006541)” and “galactose metabolism (ko00052)”; clusters III and XI shared the “starch and sucrose metabolism (ko00500)” pathway; clusters VII and VIII shared “DNA replication (ko03030)” and “mismatch repair (ko03430)” pathways.

### High-throughput small RNA sequencing identified 369 novel miRNAs

Sequencing of 30 small RNA libraries generated a total of 451.77 million clean reads after filtering and trimming, with no less than 10.98 million reads from each sample. A total of 521 miRNAs, including 152 (29.17%) known and 369 (70.83%) novel miRNAs, were identified, of which 309 belonged to 94 conserved families. The MIR167_1 family, including 45 members, was the largest, followed by MIR396 and MIR399 families (Table S5). The length of the miRNAs identified ranged from 19 to 25 nt, and the 24 nt (51.2%) miRNAs were found in the highest proportion (Figure 4A). Among the known miRNAs, 21 nt long ones accounted for the most significant proportion (73.68%), and a majority (78.57%) were characterized by a 5’ uridine residue at the first base. Among the novel miRNAs, 24 nt long ones accounted for the most significant proportion (67.75%), and a majority of them (79.60%) had a 5’ adenosine residue at the first base (Figure 4B). The average GC content of the miRNAs was 49.55%.

**Figure 4.**
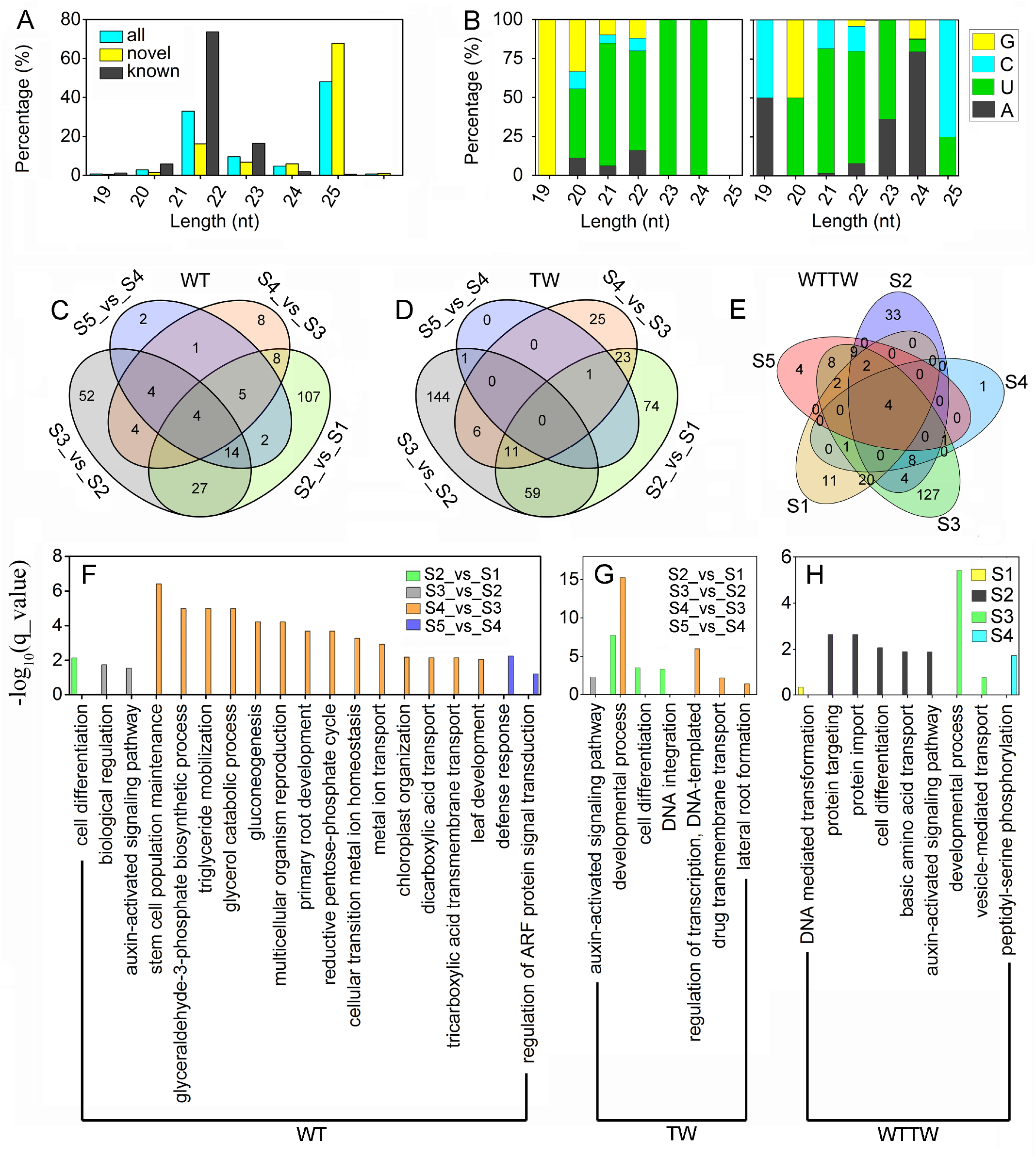
Known and novel miRNAs identified via transcriptome and small RNA sequencing using the underground thickening shoot samples of wild type (WT) Moso bamboo (*Phyllostachys edulis*) and a thick wall (TW) variant (*P. edulis* cv. Pachyloen). A and B, Length and base bias of the miRNAs. C–E, Venn diagram of differentially expressed miRNAs (DEmiRs) of the WT (C), TW (D), and WTTW groups (E). F–H, KEGG pathway analysis of DEmiRs identified from the different combinations of the WT (F), TW (G), and WTTW (H) groups.

A total of 5,790 target genes, including 5,519 (95.32%) functionally annotated ones, were predicted for 481 (152 known and 329 novel ones) miRNAs (Table S5); among them, 261 miRNAs and 453 predicted TF targets composed 2,058 miRNA-TF pairs (Table S6). The family MIR167_1 was engaged with most pairs (431), followed by MIR396 (378) and MIR159 (174).

### Shared GO terms for the targets of DEmiRs were identified in different groups

A total of 401 (76.97%) miRNAs were differentially expressed in at least one of the 13 pairwise combinations from the WT, TW, and WTTW groups; 45, 345, and 235 miRNAs showed differential expression in the WT, TW, and WTTW groups, respectively (Table S5). Among them, 169 and 243 showed stage-specific differential expression in the WT and TW groups, respectively, and 176 showed genotype- and stage-specific differential expression in the WTTW group (Figure 4C–E). The enrichment analysis found shared GO terms in the WT, TW and WTTW groups (Figure 4F–H). For example, targets of DEmiRs in WTS2_vs_WTS1, TWS2_vs_TWS1 and WTS2_vs_TWS2 shared the GO term “cell differentiation (GO:0030154)”.

### Most miRNAs were expressed in a stage- and/or genotype-specific pattern

Detailed analysis revealed that 503 (96.55%) and 515 (98. 85%) miRNAs were expressed in at least one of the samples in the WT and TW groups, respectively, while 465 (89.25%) and 464 (89.06%) miRNAs were expressed throughout the five stages. Meanwhile, 16 (3.07%) and 22 (4.22%) miRNAs were explicitly expressed in the WT and TW libraries. Cluster analysis grouped the miRNAs into two categories in the horizontal and vertical directions (Figure S3). The miRNAs were clustered into two groups according to the developmental stages rather than species in the horizontal direction; S1 and S2 were clustered into one group, while S3, S4, and S5 were clustered into the other. In the vertical direction, miRNAs in one cluster had high expression levels in WT and TW at five stages, and those in the other group were not highly expressed in all stages.

Specifically, the MIR166 family members were highly expressed throughout all stages in the WT and TW libraries (Figure 5A). Meanwhile, the MIR396 family members with a few novel ones were highly expressed at S1 and S2 and slightly expressed at S3, S4, and S5 (Figure 5B). An opposite trend was observed for miR390 and a few novel miRNAs (Figure 5C).

**Figure 5.**
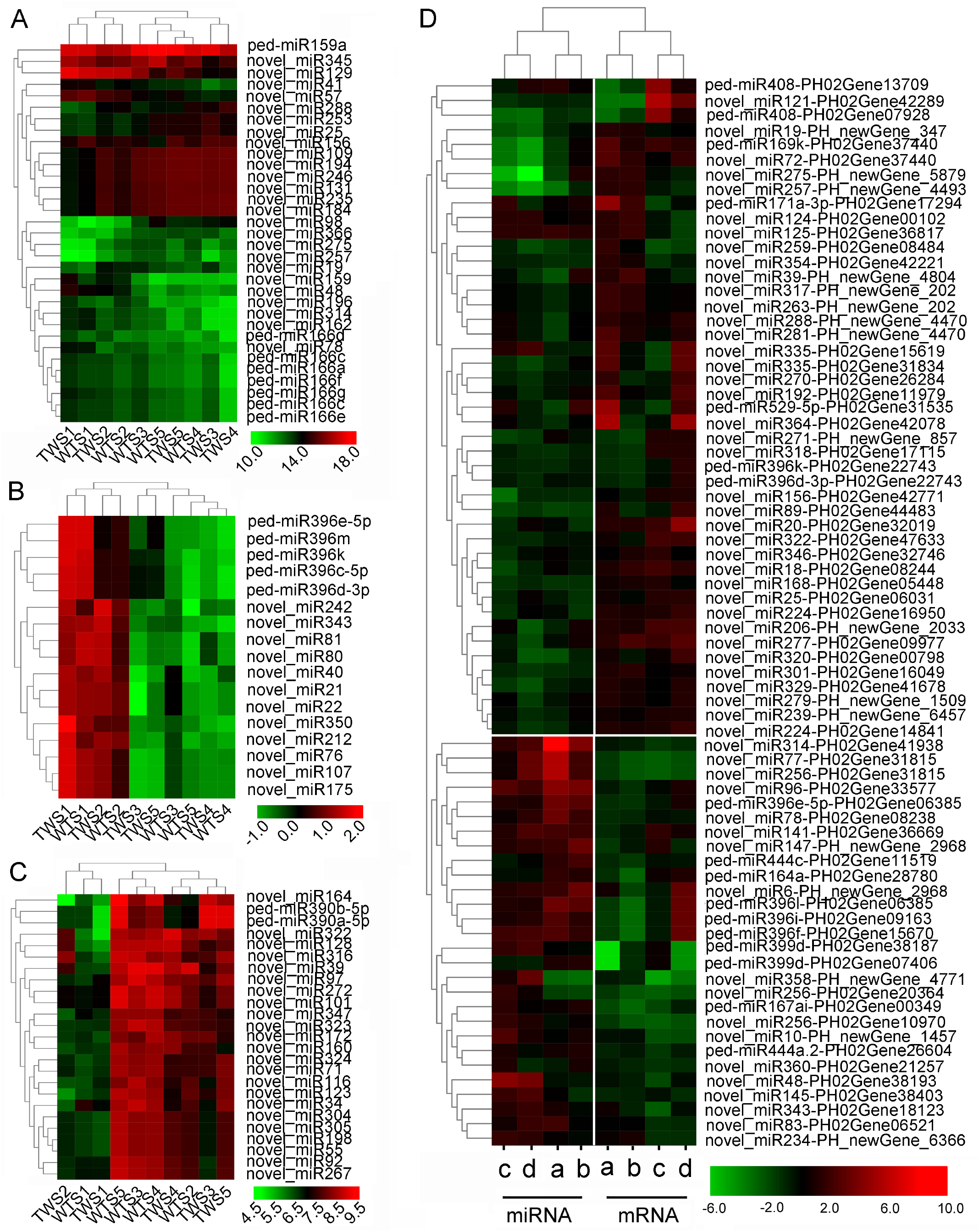
Expression patterns of miRNAs and their targets in eight pairwise combinations between two successive stages of the WT and the TW groups. A–C, Expression patterns of the miRNAs from the MIR166, MIR396 and MIR390 families. D, Expression patterns of miRNAs and their targets coherent in four dominant combinations, namely TWS2_vs_TWS3 (a), TWS3_vs_ WTS3 (b), TWS1_vs_TWS2 (c), and WT1_vs_WTS2 (d).

### Most coherently expressed DEmiRs and their predicted targets were primarily identified between the stages of WTS1-S3/TWS1-S3

Further, the correlation between DEmiRs and DEGs was analyzed. A total of 1,959 distinct miRNA-mRNA interaction pairs, including 323 in the WT group, 671 in the WTTW group, and 1,558 in the TW group (Table S7-1), were identified; the majority (1,046, 53.39%) were found in the four dominant combinations, namely WT2_vs_WTS1, TWS2_vs_TWS1, TWS3_vs_TWS2, and TWS3_vs_WTS3. Further analyses were carried out to identify whether these interactions were either coherent (the expression level of target mRNA is more when that of miRNA is less; the “DU” and “UD” patterns) or non-coherent (miRNA and its target mRNA have similar expression profiles) (Table S7-2) (Shkumatava et al., 2009).

Analyses revealed that 1,046 (out of 1959) miRNA-mRNA pairs, involving 214 miRNAs and 438 targets, were coherently expressed in at least one of the four dominant pairwise combinations (Figure 5D, Table S7-3). Besides, 176 coherent pairs were identified only in specific combinations, and most pairs (81.82%) were from the TW group. Notably, the miRNAs from the same family were consistently coherent with their corresponding targets. Maximum coherent pairs were found in WTS3_vs_TWS3. Additionally, a few pairs showed coherent expression in pairwise combinations between two successive stages. For example, members of MIR408, MIR399, and MIR444a families were coherently expressed with their targets in both TWS2_vs_TWS1 and TWS3_vs_TWS2. Most miRNAs shared targets from several TF families, resulting in a network (Figure 6).

**Figure 6.**
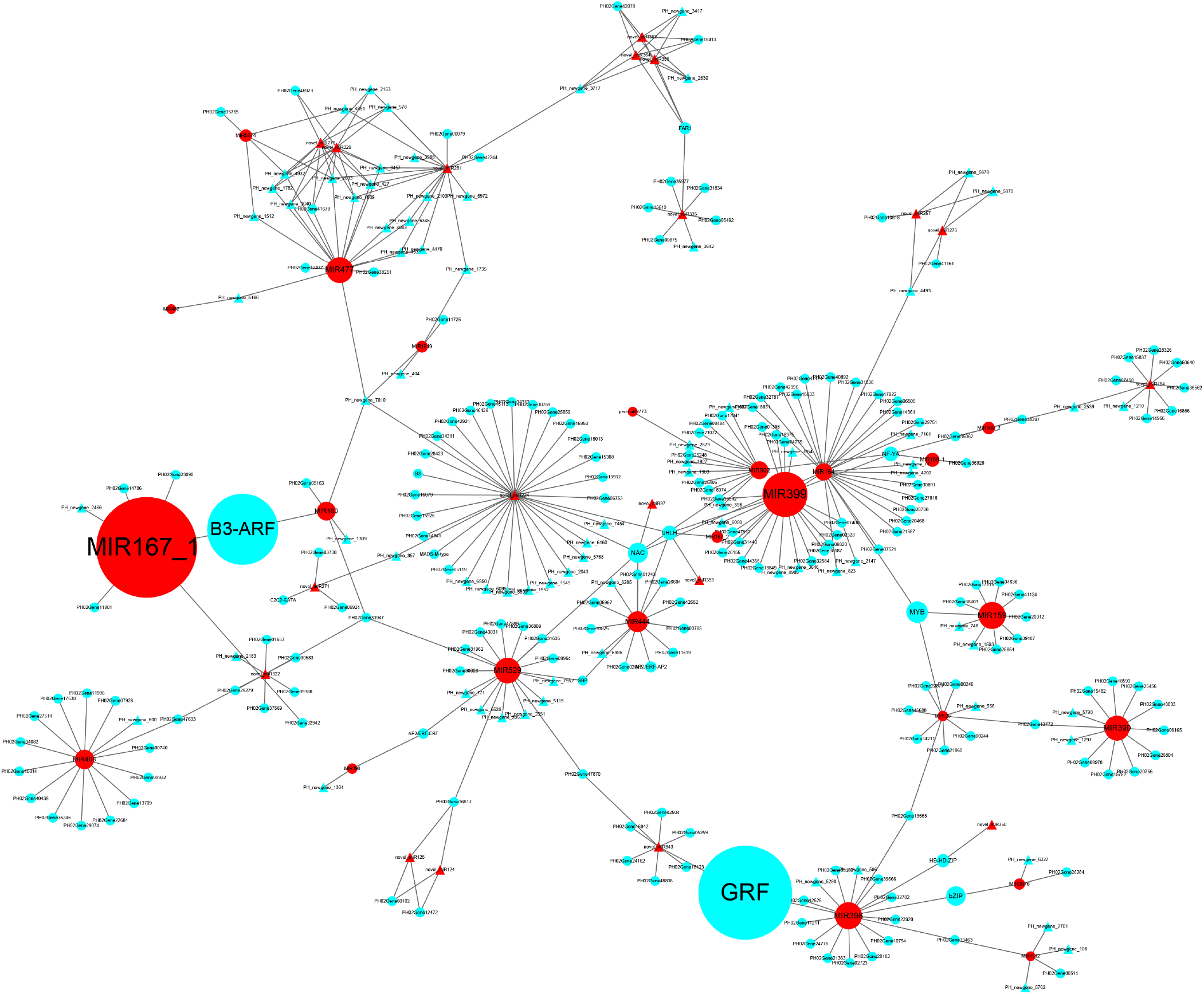
Cytoscape regulatory networks of differentially expressed miRNAs (DEmiRs) and their predicted target genes of differentially expressed genes (DEGs). Triangles represent the novel miRNAs and genes identified. Red graphics represent miRNAs, and blue graphics represent genes. The size of dots in the networks was proportional to the number of miRNAs and genes.

### Zeatin showed high concentration changes with stage

The phytohormones showed similarities between the WT and TW groups. Among the four phytohormones analyzed, changes in concentration were the largest for zeatin (ZT) (Figure 7A-B); ZT concentrations showed similar trends throughout the five stages between the WT and the TW groups. Meanwhile, the changes in gibberellic acid (GA_3_) concentration with stage were the least. At certain stages, ZT and abscisic acid (ABA) concentrations showed opposite trends.

**Figure 7.**
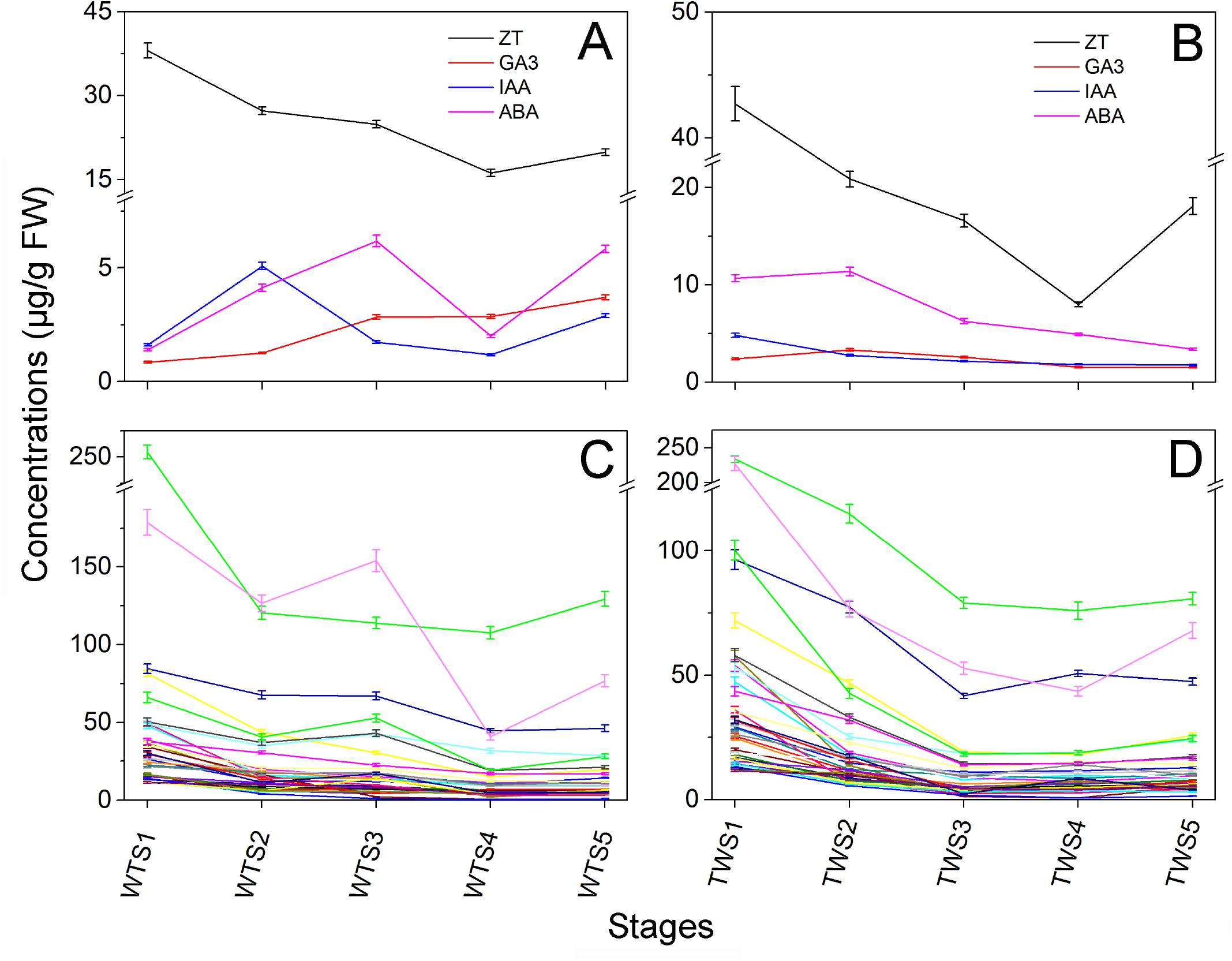
Changes in the concentrations of four phytohormones compared with the expression profiles of the predicted targets of miRNAs during five developmental stages in the wild-type (WT) Moso bamboo and the thick wall (TW) variant. A and B, Changes in the concentrations of four phytohormones in the WT (A) and the TW (B) groups. C and D, Changes in the expression profiles of 48 predicted targets for 43 miRNAs in the WT (C) and the TW (D) groups. Each point represents the mean ± SE (n = 3).

Additionally, differences in phytohormone concentrations and trends were observed between different developmental stages from the WT and the TW groups. The ZT concentration changes from S1 to S2 were opposite to that of ABA in the TW group, while it was such opposite trends in ZT and ABA from S1 to S3 in the WT group; ABA concentrations in TW were generally higher than those in WT. Meanwhile, GA_3_ concentration increased throughout the stages in WT, while it decreased gradually after reaching the highest level at S2 in TW. Notably, the expression profiles of 35 out of the 438 predicted targets identified from 1,046 miRNA-mRNA pairs throughout five stages were consistent with ZT concentrations; these targets showed similarities between TW and WT (Figure 7C–D). These genes encoded proteins, including bZIP, TF, cytochrome P450, soluble starch synthase, PK, F-box protein, and phosphoinositide phosphatase. Another 13 genes showed expression profiles different from ZT concentrations throughout five stages (Table S7-4).

### Degradome sequencing confirmed 590 miRNA-mRNA pairs

After successive filtering steps, a total of 111.89 million clean tags were generated from two distinct mixed sample pools collected from the WT and TW groups, which were processed to identify the cleavage sites. In total, 118 cleavage sites were identified for 590 miRNA-mRNA pairs, involving 126 miRNAs and 117 targets, and 101 and 109 cleavage sites were identified from the WT and TW groups, respectively. A total of 99 miRNA-mRNA pairs were from the 1,046 pairs.

Notably, *PH02Gene00233*, an auxin response factor (ARF) 13-like gene, showed two different cleavage sites for MIR160 and MIR530 families in the WT group, while no cleavage site was detected for it in the TW group. The number of cleavage sites of the same miRNA family was different between the groups. For example, five and eight cleavage sites were identified for the MIR166-mRNA pairs in the WT and TW groups, respectively. Interestingly, a few miRNAs and miRNA families showed similarities in cleave sites of targets between two groups. Generally, miRNAs of the same family cleave the same target. For example, *PH02Gene00349*, which encodes an ARF, had the same cleavage site for 30 unique miRNA members of the MIR167-1 family (Table S8).

### Dual-luciferase reporter assay indicated that 14 miRNAs specifically bind to 12 target genes

In the present study, the dual-luciferase reporter assay indicated that 14 miRNAs from ten families, specifically bond to 12 target genes. As shown in Figure 8A–H, the luciferase activity of the miRNA mimics+mRNA-WT group was lower than that of the NC mimics+mRNA-WT group (*P* < 0.01) in the transfected cells; however, no significant difference was observed between two mutant groups. For example, the dual-luciferase reporter assay confirmed that miR166 binds to *PeHOX32* (Figure 8C).

**Figure 8.**
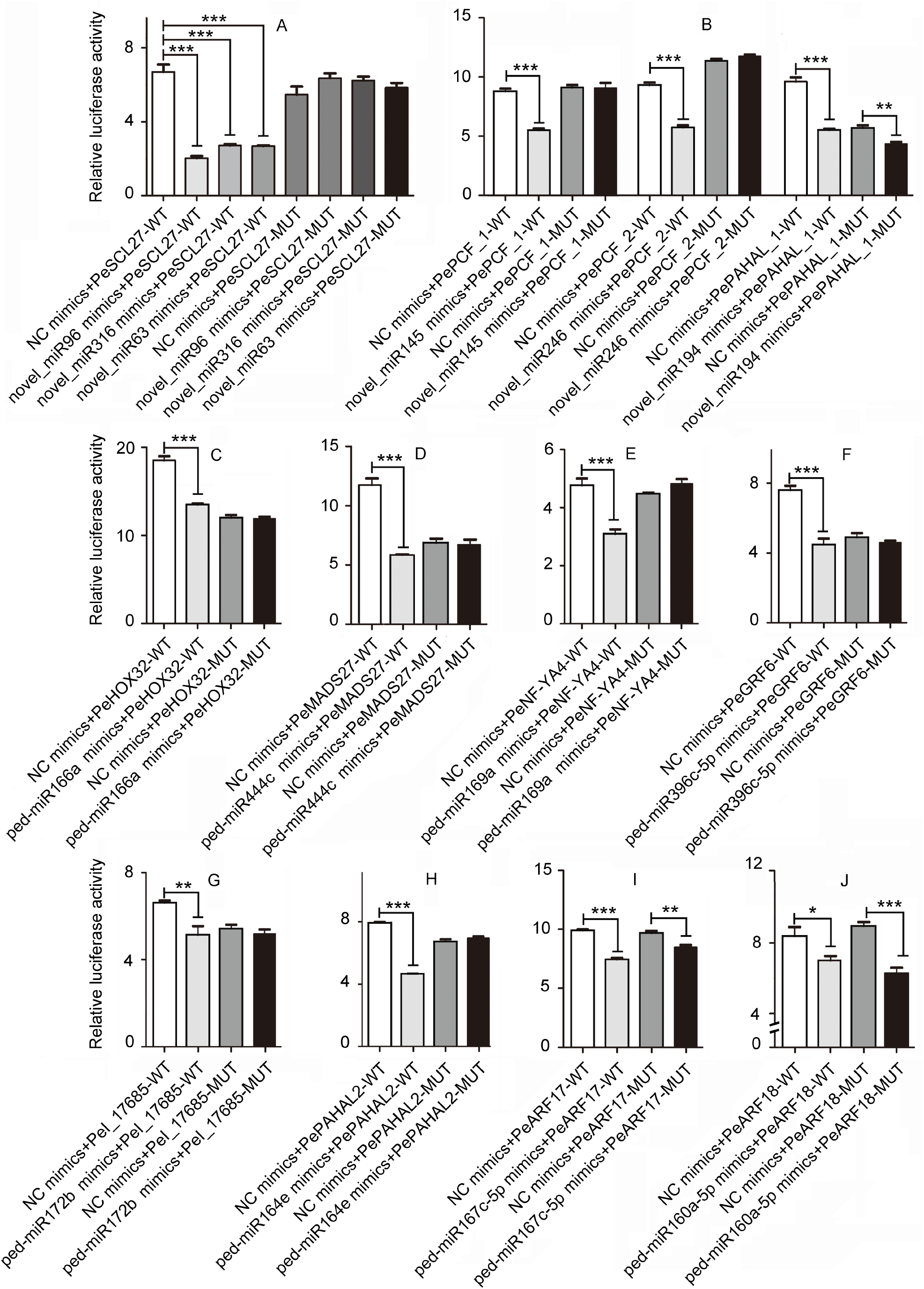
Verification of the correlation between 14 miRNAs and 12 target genes using a dual-luciferase reporter assay. 14 miRNAs were as follows: novel_miR96, novel_miR316 and novel_miR63 of the MiR171_1 family, novel_miR145, novel_miR246 and novel_miR194 of the MIR159_1 family, ped-miR166a, ped-miR444c, ped-miR169a, ped-miR396c-5p, ped-miR172b, ped-miR164e, ped-miR167c-5p, ped-miR160a-5p. 12 target genes were as follows: *PeSCL27* (*PH02Gene47151*), *PePCF6_1–2* (*PH02Gene09639* and *PH02Gene36822*), *PePAHAL_1–2* (*PH02Gene17565* and *PH02Gene15869*), *PeHOX32* (*PH02Gene31932*), *PeMADS27* (*PH02Gene21974*), *PeNF-YA4* (*PH02Gene45106*), *PeGRF6* (*PH02Gene06203*), *PeOsI* (*PH02Gene07682*), *PeARF17* (*PH02Gene03160*) and *PeARF18* (*PH02Gene37045*). Error bars indicate standard deviation (SD) of three technical repeats. Data are shown as means ± SE, n = 3. Statistical analyses were performed by *t*-test. \**P* < 0.05; \*\**P <* 0.01; *** *P* < 0.001.

Interestingly, three novel miRNAs of the MIR171_1 family, bind to *PH02Gene47151*, a putative bamboo scarecrow-like ortholog of *Brachypodium distachyon SCL27*, named *PeSCL27* (Figure 11A). And another three novel members of the MIR159 family bind to two putative bamboo TF *PePCF6s* and a *PePAHAL*, respectively (Figure 8B). Notably, as for ped-miR167c-5p of the MIR167 family and ped-miR160a-5p of the MIR160 family, significant difference of the luciferase activity was also observed between the mutant groups (Figure 8I and J). This confirmed that they bind to the corresponding targets at more than one sites.

### Fluorescence in situ hybridization (FISH) localized miR166 and miR160 in the shoot apical meristem (SAM) and the procambium

There is difference in fluorescence intensity between various tissues of the bamboo shoot in the merged image (Figure 9A). Fluorescence fluorescence intensity of miR166 is higher than that of 4,6-diamidino-2-phenyiindole 2 hci (DAPI) staining in the shoot apical meristem (SAM) and the procambium (Figure B-C). This confirmed that ped-miR166a was specifically expressed in in the shoot apical meristem and the procambium. In the same way, FISH localized novel_miR19 of the MIR166 family in the procambium and the sheath (Figure 9D-F); and FISH localized ped-miR160a-5p in the procambium and the sheath. (Figure 9G-I).

**Figure 9.**
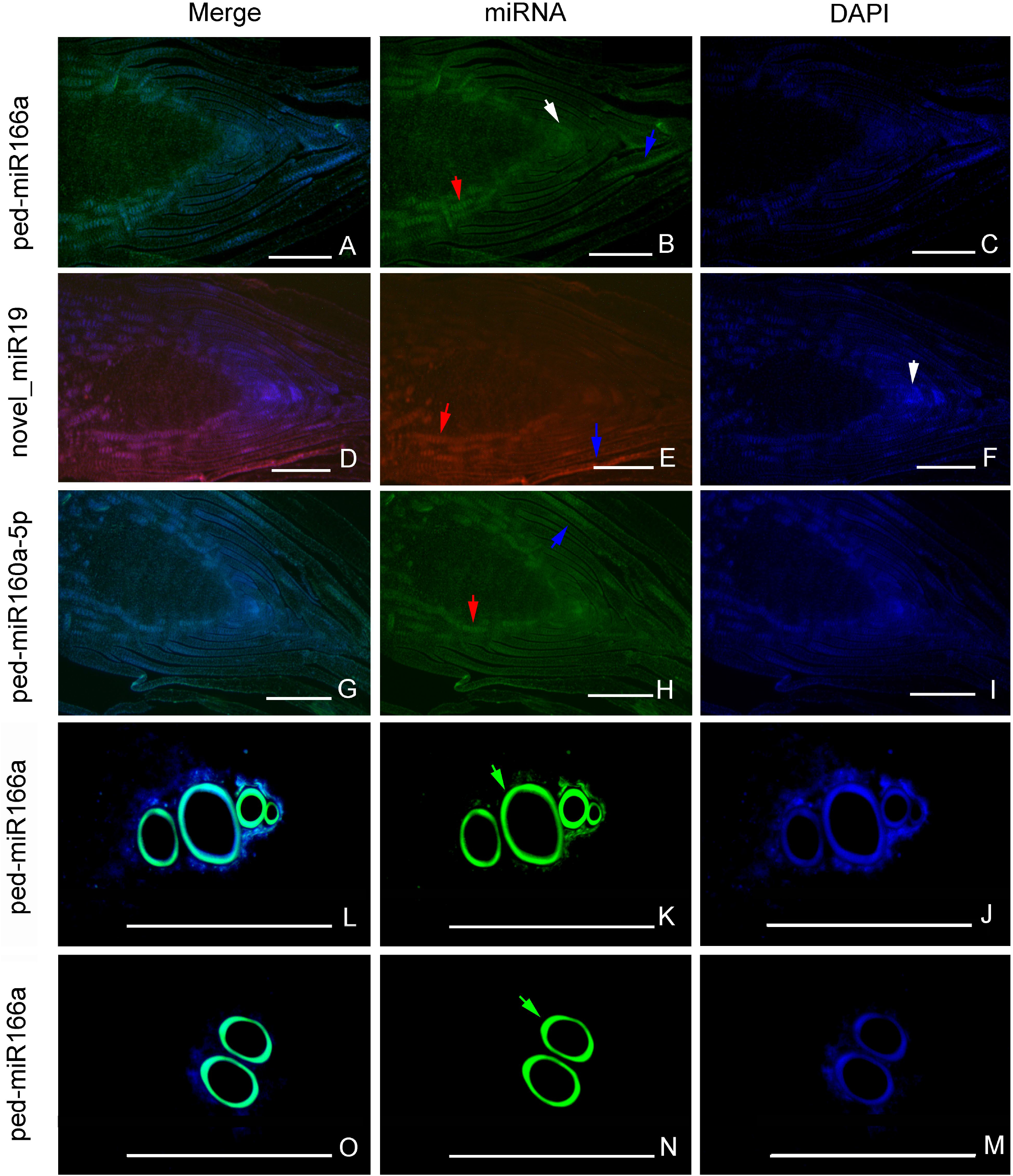
Verification of miR166 and miR160 location in the Moso bamboo shoots using fluorescence in situ hybridization (FISH). FISH localized ped-miR166a in the shoot apical meristem (white arrow) and the procambium (red arrow) (A-C), novel_miR19 of the MIR166 family in the procambium and the sheath (blue arrow) (D-F), and ped-miR160a-5p in the procambium and the sheath (G-I) of the Moso bamboo shoots at the S1 stage. FISH localized ped-miR166a in two vessels of the TW (J-L) and the WT shoots (M-O) at the S3 stage. Bar = 100 μm.

## Discussion

The present study assessed the differences in profiles and levels of genes, miRNA, and key phytohormones between the WT and TW bamboo across five developmental stages and delineated the connections among these factors in the thickening bamboo shoots.

### TFs, miRNAs, and miRNA-TF-phytohormone regulatory networks as key players during primary thickening

TFs are regulators and they carry information, which constantly updates plant development programs based on the modifications in environmental conditions, currently available resources, and sites of active organogenesis (Lai et al., 2020; Lanctot and Nemhauser, 2020. They participate in various processes, including development, immune and stress responses, and light and hormonal signaling (Schmitz and Theres, 2005). In the present study, nine TF family genes were found in high but varying proportions in the WT and TW groups. The diverse regulation modes provide a balance between the plasticity needed to rapidly adapt to the new environment and the continuity required to maintain the development of the tissues. Therefore, the differences in the ratio of the nine TF family members observed between the groups in this study indicate the changes in regulatory function, eventually causing morphological innovation of the mature culm’s wall thickness/pith diameter during evolution. Post-translational modification is a major mechanism in which miRNAs repress TFs; TFs work synergistically or antagonistically to regulate the downstream various targets. Importantly, miRNAs as parts of the miRNA-TF-plant hormone regulatory network enable complex processes with distinct developmental outcomes. For example, miRNA167-ARF regulates tissue and organ growth and development in various species (Li et al., 2020). Some similarities and differences were observed among the different families in different comparison combinations. In the present study, the expression patterns of miRNA-mRNA modules were not exactly same among three dominant pairwise combinations, indicating similarity and specificity in their regulatory functions. Meanwhile, the dual-luciferase reporter assay confirmed that miR160 and miR167_1 bind to *PeARFs*, and miR396 binds to *PeGRF*; the various members of the same miRNA family (including novel miRNAs) could degrade the same targets, and a few miRNAs had multiple target gene cleavage sites. Thus, the dissimilarity in the expression patterns of miRNA families possibly indicates the complexity in the miRNA-mRNA regulatory network, while the high similarity in the expression patterns of miRNAs from the same family ensures stability.

Hormones also play essential roles in tissue differentiation and development. In the present study, the concentration of ZT was higher than that of the other three phytohormones. A high proportion of ZT concentrations probably contributed to tissue differentiation. ZT concentration was higher than that of GA_3_, which stimulates cell proliferation and elongation (Liu et al., 2021), and modulates plant architecture (Chen et al., 2020; Miao et al., 2020) and stem elongation (Lu et al., 2020), assisting underground shoots in performing rapid internode elongation. Meanwhile, concentrations of ABA, a growth inhibitor, in the active shoots were lower than the dormant ones (Table S9), indicating that ABA could inhibits cell division and elongation of shoot tissues.

Besides, the profiles of a few predicted target genes showed similarity with ZT concentration; these genes, encoding key proteins such as cytochrome P450, bZIP TF, PK, phosphoinositide phosphatase, and F-box protein, were probably related to the synthesis and metabolism of phytohormones. Furthermore, most of miRNAs that targeted them are novel and belong to conserved families. The miRNAs, which are important for the appropriate expression of TFs, probably mediated the plant hormone response and contributed to distinct developmental outcomes in the underground shoots. The study unraveled the importance of phytohormones, especially ZT for the cell division and tissue differentiation, and emphasizes the role of miRNAs in the primary thickening bamboo shoot.

### Nonsynchronous tissue differentiation before S3 contributes to morphological differences between WT and TW

Several genes were identified from the four pairwise combinations in both WT and TW groups, indicating that the primary thickening is a continuous developmental process. In the present study, we analyzed expressions of miRNAs and genes in 13 pairwise combinations, with maximum DEmiRs and DEGs were identified in WT2/TW2_vs_WT1/TW1. Further, based on the expression patterns of 521 miRNAs, the WT1-2 and TW1-2 stages were clustered into one subgroup, and many coherently expressed miRNA-target pairs were identified from the WT1-2/TW1-2 stages. In bamboo, rapid development starts during the transition from S1 to S2, and the cross-sectional area of the active apical meristem expands dramatically (Wei et al., 2017). Various tissues, such as the pith and rib meristems, develop actively at S2. Therefore, the DEmiRs and DEGs identified in S2_vs_S1 might be involved in initiating the development of the region surrounding the meristem for both the WT and TW variant.

TW Moso, the only stable Moso bamboo variant, is characterized by a narrow pith cavity, increased vascular bundles, and a fasciated stem. The deviation in development between WT and TW initially occurs at the S1 stage; and at this stage, the apical meristem of the TW produces more vascular tissue rather than differentiate into pith cells than the WT, while both of WT and TW develop continuously and actively at S2 and S3 (Wei et al., 2017). Therefore, during this period, the pith zone in the WT shoots was apparent, while the culm wall in the TW shoots was thicker. In the present study, a total of 436 DEGs and 40 DEmiRs were identified at the S1 stage, respectively, between the WT and the TW variant. Additionally, the SAM in the TW shoot was flat and enlarged and had more cells but the same number of cell layers as that of the WT, which might account for the varied proportion of pith and vascular tissue differentiation. Moreover, the culms of WT and TW were spirally twisted but to different degrees. Because they emerge initially from the underground shoots, more effects were need to detect DEGs and/or DEmiRs in SAMs, which would probably regulated the shoot twist in the WT and the TW mutant.

Furthermore, maximum DEGs and DEmiRs were identified in WTS3_vs_TWS3 among the five relative pairwise combinations in the WTTW group. The expression profile of genes identified in the WT and TW groups mostly showed a breakpoint at the S3 stage, but in opposite directions; many miRNA-mRNA pairs were identified from the relative pairwise combinations of the S3 stage. Besides, ZT concentration decreased from S1 to S3. These indicate that it is at S3 that the cells in the WT and the TW would face distinct fates driven by expressions of DEGs and DEmiRs. The pith cavity and the wall possibly develop asynchronously between the WT and the TW variant during primary thickening. Therefore, the genotype-specific DEGs, especially those identified at S3, may play a decisive role in the differences in the mature culm’s wall thickness/pith diameter between the WT and the TW. Interestingly, no miRNA-target pairs were identified from stages later than S3, consistent with Wei et al. (2017), who demonstrated fewer apical meristematic cells of bamboo shoots at S3. These observations indicate that the SAM activity might have declined at S3, and the developmental activity might have dropped as the number of cells plummeted. However, detailed experiments are needed to verify the role of the genes in regulating spatiotemporal tissue development and differentiation and elucidate the molecular mechanism underlying primary thickening.

### Novel miRNAs might promote specific morphological characteristics in the thick wall variant

Previously, Peng et al. (2013) identified 225 miRNAs from vegetative and flowering bamboo tissues. The present study identified 521 miRNAs, with many novel ones (70.83%), substantially higher than that reported in many species, including chickpea (Garg et al., 2018; Kudapa et al., 2018), soybean (Subramanian et al., 2008), and cotton (Wang et al., 2012). Additionally, half of the coherent pairs involved novel miRNAs and were differentially expressed in combinations between two successive stages and two relative genotypes, revealing the importance of novel miRNAs for shoots in the underground developmental processes. Among them, a few miRNAs were clustered with the conserved miRNA families, such as MIR159 and MIR166. Meanwhile, many novel ones were clustered with the non-conserved and recently evolved miRNAs, such as miR7506 and miR6023. This observation suggests complex regulatory roles of miRNAs in the primary thickening growth of underground shoots.

Changes in miRNA folding, expression, processing efficiency, and size lead to changes in miRNA gene loci and produce young miRNAs (Cuperus et al., 2011). The high percentage of (59%) transposable elements in the Moso bamboo genome (Peng et al., 2013) seem to have contributed to the generation of species-specific miRNA genes (Nozawa et al., 2012). Earlier, Tang (2010) demonstrated that a few miRNA genes might evolve from target genes via coevolution, during which the species and number of miRNA families and their corresponding miRNA targets increased. Meanwhile, Peng et al. (2013) claimed that many gene duplicates involved in bamboo shoot development were generated after a whole-genome duplication event 7–12 million years ago. Some duplicates might be targets of the novel miRNA genes identified in this study. In the present study, many new genes, including TF genes, were identified, which may be responsible for the changes affecting the biochemical properties of the TF protein or its expression pattern (Lai et al., 2020). Besides, a few coherent pairs composed of novel miRNAs and new genes were also identified. Probably, the duplication, inversion, mutation, and other types of genetic drifts from protein-coding genes might have acted as the primary events in the genesis and evolution of the novel miRNA genes. Subsequent co-evolution of the miRNAs and their target genes probably ensured the maintenance and the fine-tuning of a dynamic regulatory network in bamboo.

Bamboo shoots develop from underground vegetative buds located at the nodes of a bamboo rhizome. Thus, the morphological development of the underground SAM is neither like the seedlings of *Arabidopsis* and other herbaceous plants nor like the stem or branch of fruit trees. Wei et al., (2017) suggest a close association with developmental activity of SAM cells and had found that the apical meristem of the bamboo shoot is different from that of rice (*Oryza sativa*) and maize (*Zea mays*). Meanwhile, shoots of both the WT and the TW variant appeared spirally twisted from the primary thickening growth, which is different from other monocots herbs but similar to few perennial woody trees. In the present study, several novel members and eight known ones from the MIR166 family were highly expressed; the dual-luciferase reporter assay indicated that miR166 targeted and suppressed the expression of *PeHD-ZIP*. Besides, FISH localized ped-miR166a in the shoot apical meristem and the procambium, novel_miR19 in the procambium of the Moso bamboo at the S1 stage. Previous studies indicated that the evolved or evolving novel miRNAs might be important for species-specific gene regulatory functions (Lelandais-Briere et al., 2009). Possibly, novel members were recruited to participate in the tissue differentiation to produce specific morphological characteristics in the Moso bamboo, consistent with D’Ario (2017), who demonstrated that evolution occurred via the same pathways that achieved different developmental processes.

## Conclusions

The study identified changes in the levels and profiles of DEGs, DEmiRs, their predicted targets, and phytohormones throughout the five stages of tissue differentiation. These findings indicated that the differences in miRNAome and transcriptome are essential for phenotypic plasticity, and the phytohormone ZT plays a significant role in tissue differentiation in bamboo. Furthermore, the similarities and dissimilarities in genes, miRNAs, and phytohormones identified between the WT and the TW variant indicated their potential contribution to morphological differences. In the present study, only two independent libraries from the WT and TW groups were established for degradome sequencing, and therefore, degradation sites could not be efficiently and adequately detected. Importantly, accurate sampling is required for tissues other than the whole shoots under multiple stages, which will help establish profiles of genes, miRNAs, and targets with high resolution and precision. Additional methods and efforts are required to verify the interaction between miRNAs and their predicted targets. To conclude, the study proposes candidates for bamboo genetics and breeding and the study’s findings unravel a complex association between genes, miRNAs, and phytohormones in the primary thickening of bamboo shoots.

## Materials and methods

### Plant materials

The studied plant species is Moso bamboo (*Phyllostachys edulis* (Carr.) H. de Lehaie). The wild-type (WT) Moso and the thick wall variant (TW, *P. edulis* cv. *Pachyloen*) were used for a comparative analyses; and they both grow in the bamboo garden of Jiangxi academy of forestry, Nanchang City, Jiangxi, China (28°44′35.36″ N, 115°49′36.12″ E). We collected underground shoots at stage S2-S6 defined by Wei et al., (2017) from rhizomes of three-year-old bamboo plants. Additionally, for convenience, S2-S6 stages of the WT Moso and the TW variant were named WTS1–5 and TWS1–5. After being harvested, they were immediately frozen in liquid nitrogen and stored at -80 °C until use.

To confirm the miRNA location, we collected Moso bamboo shoots at the WTS1 stages from rhizomes of three-year-old pot-seedlings that were maintained in an air-conditioned greenhouse; they were located in the National State Forestry and Grassland Administration Key Open Laboratory on the Science and Technology of Bamboo and Rattan, Beijing, CHINA. Plants were watered three times a week and maintained at 25 ± 1 °C and 50-55 % relative humidity under a 12/12 h (light/dark) photoperiod with light intensity of 1,250 μmol m^−2^ s^−1^. Besides, we also collected underground shoots from rhizomes of three-year-old bamboo plants of the WT and the TW at the S3 stage (WTS3 and TWS3); and they grow in the bamboo garden of Jiangxi academy of forestry.

### Collection of physiological and biochemical data

A total of four representative phytohormones, including zeatin (ZT), gibberellic acid (GA_3_), indole-3-acetic acid (IAA), and abscisic acid (ABA), were evaluated. ZT, GA_3_, IAA, and ABA were purchased from Shanghai Yuanye Bio-Technology Co., Ltd. (Shanghai, China). Three biological replicates (three independent shoots) were maintained for the measurement of each phytohormone, and at least three technical replicates were analyzed for each biological replicate.

Plant samples were snipped and ground to a fine powder with liquid nitrogen. Subsequently, 0.2 g of the powdered plant material was extracted with 1 mL ice-cold extracting solution (methanol: acetic acid: deionized water = 12:3:5, v/v/v) at 4 °C for 12 h. The supernatant was collected after centrifugation at 4 °C, while the residue was extracted with 0.5 mL extracting solution for 2 h, and the supernatant was collected again after centrifugation. The residue was added to 0.5 mL of the extracting solution, and the supernatant solution was extracted by ultrasonication in an ice bath for 20 min. The supernatant was collected, mixed, and evaporated in a vacuum at 40 °C until the organic phase disappeared. The aqueous residue was adjusted to 1.0 mL with petroleum ether for extraction and decolorization. The upper phase was discarded, and the lower phase was evaporated in a vacuum at 40 °C. The evaporated solid was dissolved in 0.5 mL of the mobile phase solvent, and the solution was filtered using a needle filter into a chromatographic sample bottle for testing.

The testing was performed on an LC-100 HPLC system (WUFENG, Shanghai, CHINA). Chromatographic separation was achieved on a HiQ-Sil™ C18 column (250mm × 4.6 mm, 5 μm). The four phytohormones were eluted with methanol/water (45:55, v/v) containing 0.6% acetic acid. MS/MS conditions were as follows: injection volume, 10 μL; flowrate, 0.8 mL·min^−1^; column temperature, 35 °C, and wavelength, 254 nm.

### Total RNA isolation, RNA quantification and qualification

Total RNA was isolated using miRcute Plant miRNA Isolation Kit (DP504) (TIANGEN Biotech (Beijing) CO., Ltd, Beijing, CHINA) according to the manufacturer’s instructions. RNA concentration was measured using NanoDrop 8000 (Thermo Fisher Scientific, WA, USA). RNA integrity was assessed using the RNA Nano 6000 Assay Kit of the Agilent Bioanalyzer 2100 system (Agilent Technologies, CA, USA).

### Transcriptome sequencing

#### Library preparation and sequencing

A total amount of 1 μg RNA per sample was used as input material for the RNA sample preparations. Sequencing library construction using VAHTS™ mRNA-seq V3 Library Prep Kit for Illumina^®^ (Vazyme Biotech Co., Ltd, Nanjing, CHINA) according to manufacturer’s instructions. The clustering of the index-coded samples was performed on a cBot Cluster Generation System using TruSeq PE Cluster Kit v3-cBot-HS (Illumia) according to the manufacturer’s instructions. After cluster generation, the library preparations were sequenced on an Illumina Hiseq X-TEN platform and paired-end reads were generated.

#### Data analysis

The raw reads were processed to remove primer/adaptor contamination and low-quality reads (>20% bases <Q30, and N base >10%) using Trimmomatic v0.35 (Bolger et al., 2014). The filtered reads were mapped on the Moso bamboo reference genome v2.0 using HISAT2 (http://ccb.jhu.edu/software/hisat2/index.shtml) (Kim et al., 2015). The mapped reads from each sample were assembled using StringTie (https://ccb.jhu.edu/software/stringtie/index.shtml) (Pertea et al., 2015) to generate reference-guided assemblies.

To annotate genes identified, alignments were performed based on the following databases: NR (ftp://ftp.ncbi.nih.gov/blast/db/), Swiss-Prot (http://www.uniprot.org/), GO (http://www.geneontology.org/), COG (http://www.ncbi.nlm.nih.gov/COG/), KEGG (http://www.genome.jp/kegg/), KOG (http://www.ncbi.nlm.nih.gov/KOG/), Pfam (http://pfam.xfam.org/) using BLAST v2.2.26 (Altschul et al., 1997).

Gene expression levels were estimated by Fragments Per Kilobase of transcript per Million fragments mapped (FPKM) (Florea et al., 2013). The formula is shown as follow:

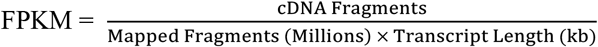

To identify differentially expressed genes (DEGs), DESeq (Wang et al., 2010) was used; genes with |log2(FC)| ≥ 1.00 and a Benjamini-Hochberg false discovery rate (FDR) corrected *P* - value < 0.01 were assigned as differentially expressed. The derived genes were represented by “A_vs_B”; for example, WT1_vs_WT2 represents differentially expressed genes between WTS1 and WTS2.

#### PPI (Protein Protein Interaction)

Sequences of the DEGs were blast (blastx) to the genome of a related species (the protein protein interaction of which exists in the STRING database: http://string-db.org/) to get the predicted PPI of these DEGs. Then the PPI of these DEGs were visualized in Cytoscape (Shannon et al., 2003).

### Small RNA sequencing

#### Library preparation and sequencing

Small RNA libraries were constructed using the Illumina TruSeq Small RNA Library Prep Kit (Illumina Inc., CA, USA) according to the manufacturer’s instructions. The clustering of the index-coded samples was performed on a cBot Cluster Generation System using TruSeq PE Cluster Kit v4-cBot-HS (Illumia) according to the manufacturer’s instructions. The library preparations were sequenced on an Illumina HiSeq X-Ten platform.

#### Data analysis

The raw reads obtained were processed to remove reads with adaptor/primer contamination and low-quality reads (>20% bases <Q30, and N base >10%); and those shorter than 18 nt and longer than 30 nt were also discarded.

#### Identification of the miRNAs

The derived reads were further screened against rRNA, tRNA, snRNA, snoRNA and other ncRNA as well as repeat sequences by mapped to various RNA database, which include Silva (http://www.arb-silva.de/), GtRNAdb (http://lowelab.ucsc.edu/GtRNAdb/), Rfam (http://rfam.xfam.org/) and Repbase (http://www.girinst.org/repbase/) by using Bowtie (Langmead et al., 2009); and the remaining unannotated reads were then mapped onto the Moso bamboo reference genome v2.0. Mapped reads from each sample were mapped onto plant miRNAs from miRBase (release 22, http://mirbase.org/) for identification of known miRNAs using Bowtie (Kozomara et al., 2019).

And the unaligned unique reads were further used for novel miRNA prediction using miRDeep-P (Yang and Li, 2011). It revolves around the secondary structure predicted using Randfold v2.0 (https://anaconda.org/bioconda/randfold), mainly including star miRNA evidence, less than six nucleotides difference between mature and star miRNA lengths, the Dicer cleavage site and the minimum free energy (Meyers et al., 2008). Further, miRNAs identified were clustered into families based on sequence similarity.

### Expression analysis of miRNAs

To quantify the abundance of miRNA, TPM (Tags Per Million reads) algorithm (Li et al., 2010) was used to normalize expressions. Here, TPM values were calculated using the following equation:

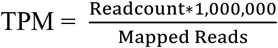

In the formula, Readcount represents the number of reads mapped onto a certain miRNA; Mapped Reads denotes the number of reads mapped onto all miRNAs in the reference genome v2.0.

To identify differentially expressed miRNAs, DESeq2 v1.6.3 (Love et al., 2014) was used, and the row z-score method was used to normalize the raw read counts of miRNA; miRNAs with |log2(FC)| ≥ 1.00 and a Benjamini-Hochberg false discovery rate (FDR) corrected *P* - value < 0.01 were assigned as differentially expressed. The derived miRNAs by this analyses was differentially expressed miRNAs (here names “DEmiRs”) and was represented by “A_vs_B”; for example, S01_vs_S02 represents the DEmiRs between S01 and S02.

Hierarchical clustering analysis was performed on the DEmiRs by using BMKCloud (www.biocloud.net), and miRNAs with the same or similar expression patterns were clustered.

#### Targets prediction and function annotation

Potential miRNA targets were identified using the TargetFinder v1.6 (Allen et al., 2005). Annotation of these sequences were performed to query against various database mentioned in “*Data analysis*”.

### Degradome library construction and analysis

The total RNA from the WT and the TW groups were pooled separately to generate two unique libraries for degradome sequencing, one representing the WT and the other the TW variant. The library was constructed as previously described (German et al. 2009) with slight modifications followed by sequencing on an Illumina Hiseq 2500 platform. The raw reads (single-end; 50 bp) were processed using Trimmomatic (v0.35) to remove low-quality reads, reads with ‘N’s, and any reads with adaptor and primer contamination. The filtered reads were searched against all other non-coding RNA sequences from Rfam except miRNA using Bowtie, followed by removing reads aligning to rRNAs, tRNAs, snoRNAs, and repeats. The clean reads obtained were mapped onto the reference genome (v2.0), allowing a maximum of one mismatch. The mapped reads were processed using the CleaveLand (v4.4) pipeline (Addo-Quaye et al., 2009) to predict the miRNA cleavage sites. The cleavage sites at the 10^th^ position relative to the aligned miRNA were considered significant at *P* ≤ 0.05.

### Fluorescence in situ hybridization (FISH)

Specific FISH probes were designed and synthesized by Abiocenter (Beijing, China). The hybridization was performed in cells and tissues of the bamboo shoots at the S1 stages. All images were analyzed on a confocal laser scanning microscope (Leica Microsystems, Mannheim, Germany). The FISH probe sequences are shown as follows: ped-miR160a-5p: 5′-TGGCATACAGGGAGCCAGGCA-3′; novel_miR19: 5′-AGGGATTGAAGCCTGGTCCGA-3′; ped-miR166a: 5′-GGGGGAATGAAGCCTGGTCCGA -3′.

### Luciferase reporter assay

Wild-type (WT) of miRNAs and mRNAs of length 200bp, including the predicted splicing sites and the flanking sequence, and mutant type (MUT) after site-directed mutation of WT target sites were synthesized artificially. Then, they were cloned into pmirGLO vector (Wuhan GeneCreate Biological Engineering Co., Ltd., Wuhan, China) between *Nhe*I and *Xho*I (TaKaRa, Nojihigashi, Japan), and the empty plasmid was transfected as the control group. After confirmation by sequencing, MUT and WT vectors were co-transfected with mimic-NC or miRNAs mimic to 293T cells, respectively. Cells were collected and processed 48 h after transfection according to the manufacturer’s protocol with the Luciferase reporter assay kit (RG005, Beyotime, Shanghai, CHINA). The outcomes were quantified in each well as the proportion of activity of firefly luciferase/Renilla luciferase. Three independent experiment was performed likewise.

### Accession numbers

Sequence data have been deposited in NCBI Sequence Read Archive (SRA, http://www.ncbi.nlm.nih.gov/sra) with the BioProject ID PRJNA753616.

## Supplemental Data

**Supplemental Figure S1**. KEGG pathway enrichment of differentially expressed genes (DEGs) identified from13 combinations of the WT, TW and WTTW groups. The WT and the TW group were composed of four pairwise combinations each between two successive stages (WTS1/TWS1_vs_WTS2/TWS2, WTS2/TWS2_vs_WTS3/TWS3, WTS3/TWS3_vs_WTS4/TWS4 and WTS4/TWS4_vs_WTS5/TWS5), and the WTTW group was composed of five pairwise combinations between two relative stages (TWS1– 5_vs_WTS1–5).

**Supplemental Figure S2**. KEGG pathway enrichment of differentially expressed genes (DEGs) identified from 13 combinations of the WT, TW and WTTW groups.

**Supplemental Figure S3**. Expression patterns of known and novel miRNAs identified in the WT, TW and WTTW groups.

**Supplemental Table S1**. Significant enriched GO terms for differentially expressed genes (DEGs) identified in five dominant pairwise combinations, namely WTS2_vs_WTS1, WTS4_vs_WTS3, TWS2_vs_TWS1, TWS3_vs_TWS2 and TWS3_vs_WTS3.

**Supplemental Table S2**. 3,543, 2,397 and 780 genes identified encode transcription factors (IFs), protein kinases (PKs) and transcription regulator (TRs), respectively.

**Supplemental Table S3**. GO terms enrichment of differentially expressed genes (DEGs) in 11 clusters.

**Supplemental Table S4**. KEGG pathway enrichment of differentially expressed genes (DEGs) in 11 clusters.

**Supplemental Table S5**. Table S5. Known and novel miRNAs identified via transcriptome and small RNA sequencing using the underground thickening shoot samples of wild type (WT) Moso bamboo (*Phyllostachys edulis*) and a thick wall (TW) variant (*P. edulis* cv. Pachyloen). **Supplemental Table S6**. 2,058 miRNA-TF gene pairs composed of differentially expressed miRNAs (DEmiRs) and their predicted targets.

**Supplemental Table S7**. Expression patterns of 1,046 miRNA-mRNA pairs coherent in four dominant combinations, namely TWS3_vs_TWS2 (A), TWS3_vs_WTS3 (B), TWS2_vs_TWS1 (C) and WT2_vs_WTS1 (D).

**Supplemental Table S8**. A total of 118 cleavage sites for 590 miRNA-mRNA pairs were confirmed by using degradome sequencing.

## Acknowledgments

This work was supported by the National Natural Science Foundation of China (NSFC) (No. 31901371) and the Special Funds for Fundamental Scientific Research on Professional Work Supported by International Center for Bamboo and Rattan (No. 1632019025).

